# Interstitial infusion of purified collagenase Clostridium histolyticum in the cirrhotic liver causes rapid reduction in fibrosis with minimal liver toxicity

**DOI:** 10.1101/2024.10.08.617298

**Authors:** Ben D. Leaker, Christiane Fuchs, Emma Wise, Joshua Tam, R. Rox Anderson

**Affiliations:** Health Sciences and Technology, Harvard-Massachusetts Institute of Technology, Cambridge, MA, USA; Wellman Center for Photomedicine, Massachusetts General Hospital, Boston, MA, USA; Department of Dermatology, Harvard Medical School, Boston, MA, USA

## Abstract

Crude mixtures of matrix-degrading enzymes called collagenase Clostridium histolyticum (CCH) have been used to efficiently breakdown tissue for many years. Recently, direct injection of purified CCH has been successfully developed as a treatment for Dupuytren’s contracture (DC), a fibrotic disorder of the hand. Given similar histologic and mechanical features between the fibrous bands in DC and cirrhosis, a similar approach may be feasible for the treatment of cirrhosis. Crude and purified CCH were first compared through composition and substrate specificity. The biodistribution of a macromolecule delivered via interstitial infusion in the liver was mapped and quantified with a fluorescent dextran tracer to design a protocol for efficient delivery throughout the liver with minimal off-target exposure. Safety and efficacy of interstitial CCH infusion in the liver was investigated in cirrhotic mice using serum markers of injury and histological analysis of fibrosis. Purified CCH showed high purity and efficient degradation of type I collagen. Tracer experiments showed that interstitial infusion in one lobe of the liver will reach the entire organ in the mouse. A significant amount of the tracer was also found to enter the bloodstream where it is cleared by the kidneys, but was not found to significantly infiltrate other organs. Mild elevation in liver enzymes AST and ALT were observed 1d after infusion of purified CCH, but this was not significantly different than infusion with saline. No elevation in creatinine was observed. Cirrhotic mice infused with purified CCH showed 38% reduction in collagen proportionate area compared to mice infused with saline. These results show interstitial delivery of purified CCH in the cirrhotic liver rapidly decreases collagen content with minimal liver toxicity. This strategy merits further study as a potential treatment for cirrhosis.

## Introduction

Cirrhosis is a significant health concern throughout the world. It is the common final stage of chronic liver diseases, including alcoholic liver disease, viral hepatitis, and metabolic dysfunction-associated steatotic liver disease (MASLD). Cirrhosis is characterized by the accumulation of collagen and remodeling of the tissue into a nodular architecture, in which dense bands of fibrotic collagen encapsulate functional liver tissue. These structural changes can be accompanied by severe loss of liver function and may ultimately cause liver failure. The only current curative treatment option for cirrhosis is liver transplant.

It is well established that the liver is capable of spontaneous recovery if the underlying injury is resolved early enough^1,2^. There has been great progress in developing drugs to treat specific liver diseases or halt progression before substantial damage has occurred. However, 40% of cases are asymptomatic until cirrhosis is established^3^ so many patients will not be identified early enough for this approach to be effective. As such, there is also a significant need for treatments targeted to patients with established cirrhosis, when spontaneous reversal no longer occurs after the underlying injury is removed.

Fibrotic disorders occur throughout the body. Compared to other fields, dermatology has favored more direct and potent interventions for the treatment of fibrosis. In recent years, direct injection of a purified collagenase clostridium histolyticum (CCH) has been successfully developed as a treatment for Dupuytren’s contracture (DC)^4^, a localized fibrotic condition in the palmar fascia. DC shares a number of features with liver fibrosis, including excess collagen production, increased collagen crosslinking, hypoxia, vascularization, and myofibroblast differentiation, proliferation and contraction^5-8^. This disease progresses from a nodule – characterized by inflammation, myofibroblast proliferation, and matrix deposition – to a cord – which is hypocellular, crosslinked, and histologically more similar to mature scar tissue^9,10^. Injection of the purified CCH causes rapid degradation of collagen which mechanically weakens the cord within just 1 day. The fingers are then manipulated to rupture the cord, which removes the pro-fibrotic mechanical stimulus^4,11^. Purified CCH injection has also been successfully used in Peyronie’s disease, a similar fibrotic condition occurring in the penis^12,13^.

CCH is a mixture of enzymes secreted by the clostridium histolyticum species of bacteria. Crude and partially purified preparations of CCH are routinely used for tissue digestion during the isolation of primary cells. Crude CCH is a poorly defined mixture of enzymes including potent collagenases and a variety of other enzymes, such as clostripain and neutral proteases, which broadly digest non-collagenous proteins as well. The commercial version of CCH used for DC is purified for two collagenase enzymes^14^. This purified mixture was reported to have strong activity against collagen types I and III^14^, the two primary excess matrix components in both DC^9,15^ and cirrhosis^16^. Furthermore, the histologic features of the DC cord – namely densely packed and highly crosslinked collagen – should make it resistant to enzymatic degradation, similar to the fibrous septa in cirrhosis^17,18^. The success of CCH treatment in DC shows that these potent enzymes can overcome this obstacle.

In this work, we aimed to determine whether direct delivery of purified CCH could be used as a treatment for cirrhosis. We first compare crude and purified CCH by molecular weight composition and substrate specificity and demonstrate the potency of purified CCH on *ex vivo* cirrhotic mouse liver tissue sections. We then characterize the biodistribution of a macromolecule delivered via interstitial infusion in the liver using a fluorescent dextran tracer. Finally, we evaluate the safety and efficacy of purified CCH delivered via interstitial infusion in the liver with a mouse model of cirrhosis.

## Methods

### Crude and Purified CCH Analysis

Molecular weight analysis of the crude (Sigma Aldrich #C5138) and purified CCH (VitaCyte #001-1050) was performed using the capillary Western total protein assay^19^ (Jess, ProteinSimple). Sample preparation was done as follows: in brief, protein was diluted to 0.03mg/mL and mixed with a fluorescent standard (EZ Standard Pack # PS-ST01EZ). Samples were loaded onto respective wells in the cartridge. Loading of capillary cartridges with 12-230 kDa separation (ProteinSimple SM-W004-1) was performed according to the manufacturer’s protocol (Total protein detection module for Chemiluminescence #DM-TP01). Run was set up using the instrument software (Compass for SW). Jess Total Protein data were analyzed using the Compass for SW software, Version 6.1.0, Build ID: 0111, Revision ID 7214faa from ProteinSimple.

To assess substrate specificity, enzyme activity was measured against type I collagen, casein, and Nα-Benzoyl-L-arginine ethyl ester (BAEE). Degradation of type I collagen was measured with the EnzChek Collagenase Assay kit (Thermo Fisher #E12055) with quenched fluorescent type I collagen (Thermo Fisher #D12060) according to manufacturer instructions. Degradation of casein was measured with the Pierce Colorimetric Protease Assay kit (Thermo Fisher #23263) according to manufacturer instructions. Tryptic activity was measured using 0.23mM BAEE (Sigma Aldrich #B4500) in sodium phosphate buffer with 2mM 2-mercaptoethanol. The rate of change in absorbance at 253nm was measured with a SpectraMax M5 plate reader (Molecular Devices).

### Animals

All animal work was approved by the Massachusetts General Hospital Institutional Animal Care and Use Committee and performed in accordance with all relevant guidelines. Male C57BL/6 mice were purchased from Charles River Laboratories. Animals were housed in a controlled environment with food and water ad libitum. To induce cirrhosis, mice received intraperitoneal injections of thioacetamide three times per week for 15 weeks. The dose escalated from 50mg/kg for the first 2 doses, 100mg/kg for doses 3-5, 200mg/kg for doses 6-10, 300mg/kg for doses 11-15, then 400mg/kg until the end of the dosing schedule as established previously.^20^ Animals underwent the infusion procedure 1 week after the final dose to allow for washout of the chemical.

### Ex vivo collagen degradation

Liver samples from cirrhotic mice were embedded in OCT, snap frozen on dry ice, and cut into 12µm sections in a cryostat. Sections were then incubated with 0.5µg/mL purified CCH in normal saline supplemented with 0.3mg/mL calcium chloride dihydrate (Sigma Aldrich #C7902). Calcium is a critical cofactor for collagenase activity. Control sections were incubated with normal saline supplemented with 0.3mg/mL calcium chloride dihydrate. After 1hr, 2hr, 3hr, 4hr, or 5hr, sections were rinsed with normal saline, fixed with fresh formaldehyde, and stained with picrosirius red as described below.

### Surgical Protocol & Intrahepatic Infusion

Animals were anesthetized with isoflurane and prepared for surgery by clipping fur on the abdomen and disinfecting with 10% povidone-iodine. A laparotomy was performed with a short midline incision beginning from the xyphoid. The medial lobe of the liver was exposed using sterile swabs. The abdomen was then covered with a sterile Tegaderm film to protect the liver and limit water loss.

For the infusion, a 28G needle was connected to a 1mL syringe with a sterile silicone tube. For biodistribution experiments the syringe was filled with sterile-filtered fluorescent dextran tracer (Thermo Fisher, #D1864 and #D1830) in normal saline at a concentration of 2mg/mL. For efficacy experiments, the syringe was filled with sterile-filtered purified CCH at varying concentration in normal saline supplemented with 0.3mg/mL calcium chloride dihydrate. Control animals for biodistribution experiments received sterile-filtered normal saline, and control animals for efficacy experiments received sterile-filtered normal saline supplemented with 0.3mg/mL calcium chloride dihydrate. The needle was inserted several millimeters into the medial lobe. Using a syringe pump, interstitial infusion was performed at a rate of 100µL/hr for 1 hour.

After the infusion, the muscle layer was closed with 5-0 absorbable sutures and the skin was closed with wound clips. For biodistribution experiments, animals were euthanized via cardiac puncture 1hr, 6hr, 1d, 2d, and 3d after the infusion. For efficacy experiments, animals were euthanized 1d and 5d after the infusion.

Blood collected at the time of euthanasia was left to coagulate at room temp for 30min-1hr, then centrifuged at 2000g for 15min to separate the serum. Serum samples were stored at -20C until sent for analysis of AST, ALT, and creatinine levels.

### Biodistribution

70kDa dextran conjugated to Texas Red (Thermo Fisher, #D1864) was used to characterize the distribution of a macromolecule delivered via interstitial infusion in the liver. Distribution through the liver during infusion was measured with an IVIS Lumina Series III *in vivo* fluorescent imaging system (Caliper Life Sciences) at 10-minute intervals. The IVIS was also used to image fluorescence of the liver, kidneys, lungs, heart, and spleen after 1hr of infusion.

Lysine-fixable 70kDa dextran conjugated to Texas Red (Thermo Fisher, #D1830) was used for histological assessment of the tracer distribution. After euthanasia, the medial and left lobes of the liver, as well as the kidneys and lungs, were embedded in OCT and snap frozen on dry ice. 10µm sections were cut in a cryostat and fixed with formalin. Slides were imaged with an epifluorescence microscope and mean fluorescence intensity was quantified in ImageJ. Tissue from untreated mice was used to establish baseline fluorescence intensity in each organ.

### Histology & Immunostaining

Tissue for histology was collected at the time of euthanasia. Samples of the medial and left lobes were fixed in formaldehyde, embedded in paraffin, cut to 5µm sections, and stained with picrosirius red. Slides were imaged with a Nanozoomer 2.0 HT slide scanner.

### Image Quantification

For fibrosis assessment, 5x magnification images across the entire section were exported from the whole-slide scans captured with the Nanozoomer. All image names were then masked and randomly ordered with the Blind Analysis tool in ImageJ. Collagen proportionate area (CPA) was calculated in ImageJ^21,22^.

### Statistical Analysis

Two tailed Student’s t-test was used to assess statistical significance with a p-value threshold of 0.05. Results are plotted as mean ± SD.

## Results

### Characterization of Purified and Crude CCH

To ensure purity of the enzyme, molecular weight analysis was performed for both crude CCH and purified CCH with the capillary Western total protein assay (Fig. 1A-C). As expected, the crude CCH preparation demonstrated multiple peaks at a variety of molecular weights. The purified CCH showed one strong, distinct peak around 116 kDa, indicating high purity. A small peak was also observed around 230 kDa, however this peak was present at a similar level across a serial dilution of the purified CCH (Supplementary Figure 1), likely indicating this is an artifact rather than impurity in the CCH preparation.

**Figure 1:**
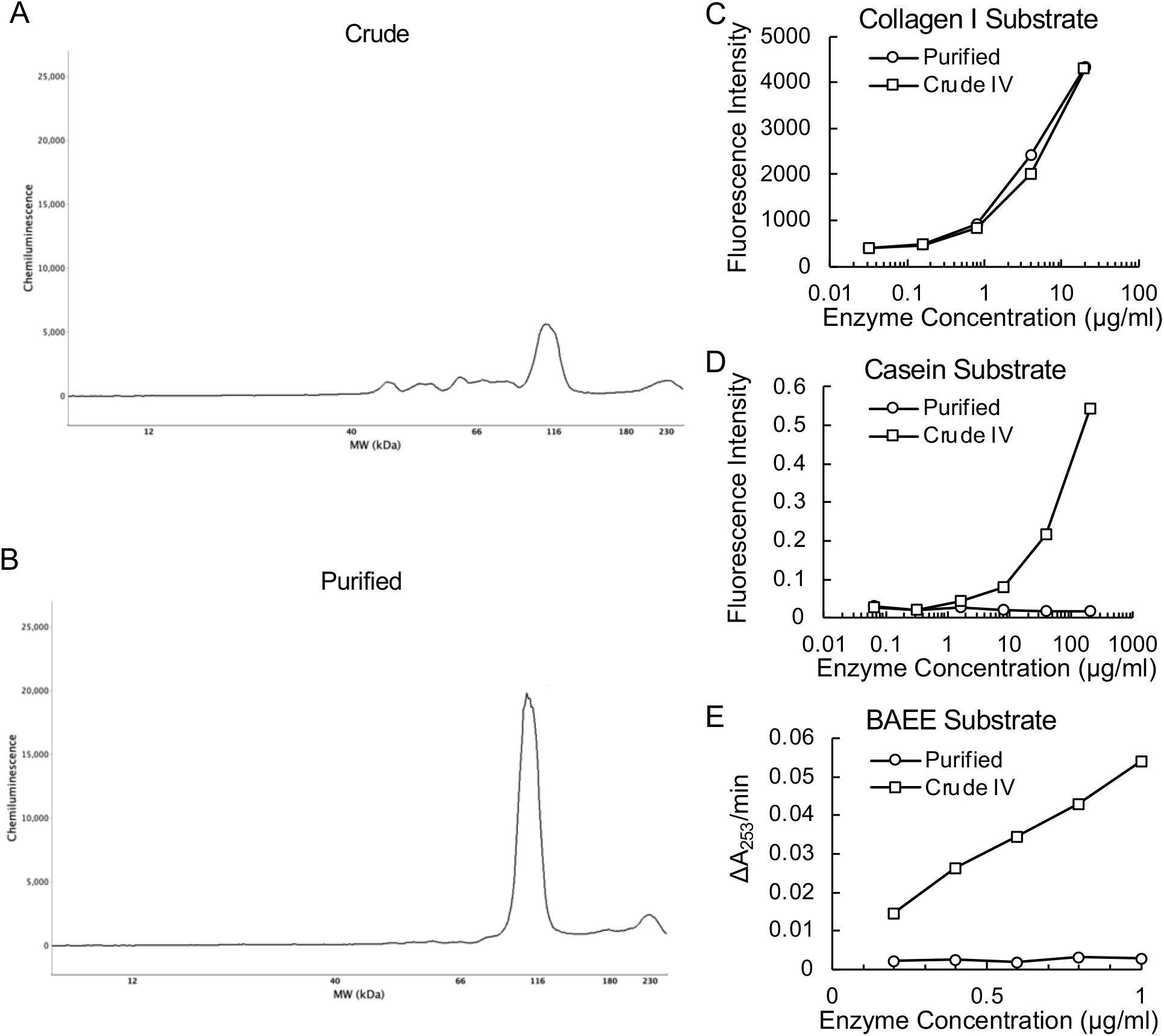
Analysis of crude and purified CCH. (A,B) Molecular weight analysis of crude and purified CCH, respectively. Crude CCH has components at a variety of molecular weights, while the purified CCH has one distinct peak around 116kDa. (C) Enzyme activity against collagen I substrate. (D) Enzyme activity against casein substrate. (E) Enzyme activity against BAEE, a synthetic substrate for measuring tryptic activity. Crude CCH degrades casein and BAEE rapidly due to the many proteases it contains. Purified CCH shows no activity against casein or BAEE, but maintains high activity against collagen I.

To further assess purity as well as substrate specificity, enzyme activity was measured against type I collagen I, casein, and BAEE (Fig. 1D,E). Crude CCH showed strong activity against all substrates, while the purified CCH only exhibited activity against collagen.

The enzyme mixtures were then tested on frozen sections of cirrhotic mice livers to quantify collagen degradation *ex vivo*. Incubating sections with crude CCH led to severe tissue dissociation, causing sections to fall apart during staining. Sections incubated with purified CCH showed no signs of tissue dissociation and had significantly reduced collagen proportionate area within 3hrs (Fig. 2).

**Figure 2:**
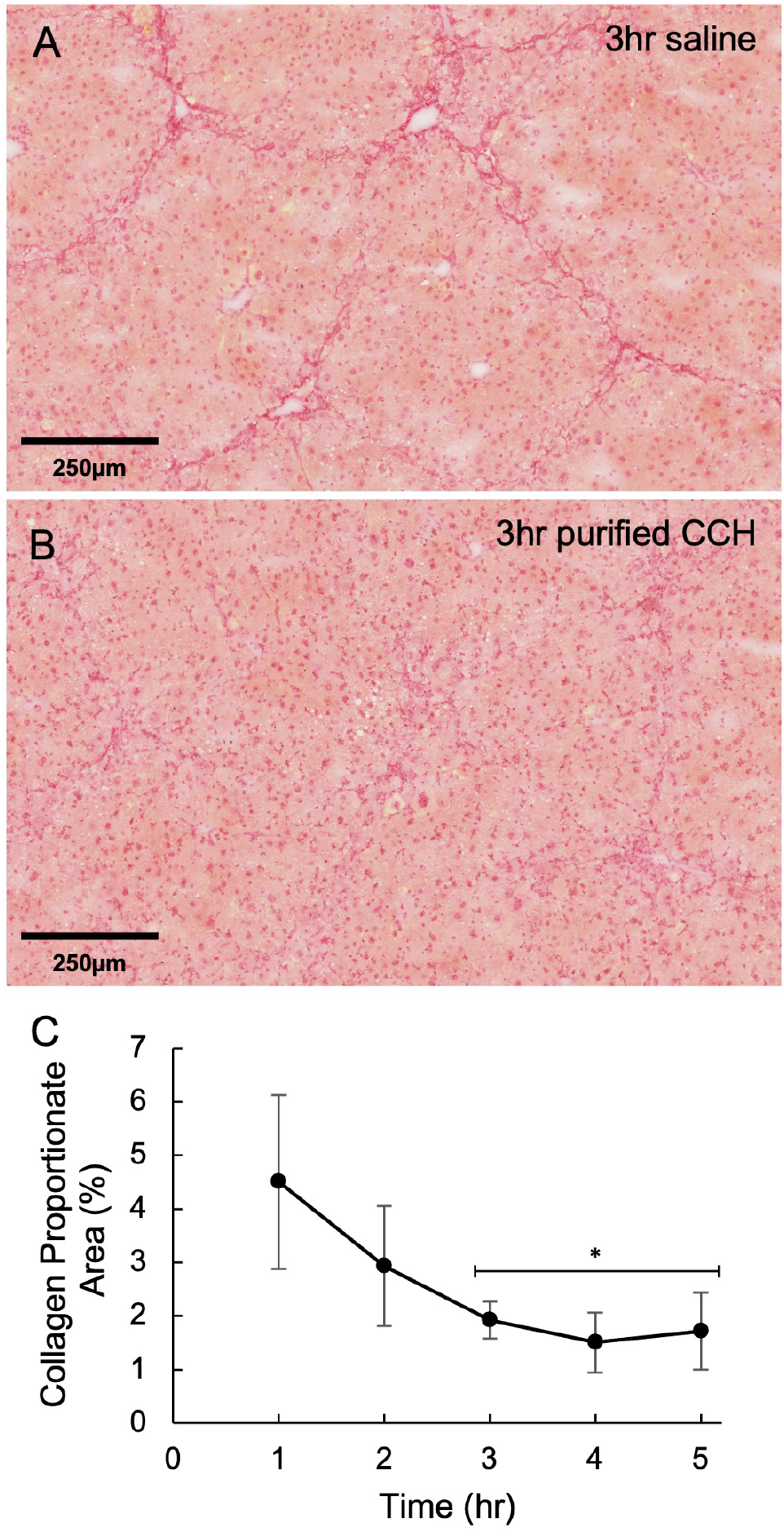
*Ex vivo* collagen degradation in cirrhotic liver sections. (A) Sections incubated with saline show distinct fibrous bands. (B) After 3 hrs with 0.5µg/mL purified CCH, the fibrous bands are nearly completely dissolved, but there is no bulk tissue dissociation as found with crude CCH. (C) Quantified reduction in collagen over time with purified CCH shows rapid degradation. Mean ± SD; n=3 per timepoint. *p<0.05 compared to saline controls.

These results verify that the purified CCH is not significantly contaminated with non-collagenase enzymes, which may cause tissue damage and cell toxicity *in vivo*, and that it is capable of rapid degradation of type I collagen without broadly disrupting tissue integrity.

### Biodistribution after Interstitial Infusion in the Liver

We next used a fluorescently labeled dextran to design an appropriate infusion protocol and characterize how the enzyme would spread after infusion. Due to autofluorescence in the cirrhotic liver, biodistribution experiments were performed in healthy mice and interlobular distribution was confirmed in the cirrhotic liver (Supplementary Figure 2).

Using live imaging, we found that the entire liver was fluorescent after 60min at an infusion rate of 0.1mL/hr (Fig. 3). We also used this system to image key organs after 1hr infusion (Fig. 4). We observed strong fluorescence throughout the liver as well as the kidneys, with no significant signal in the lungs, heart, or spleen. These results show that the entire liver receives the infused material, but a significant amount also enters the bloodstream before being cleared out by the kidneys. It should be noted that clearance of CCH through the kidneys likely differs significantly from dextran, as will be discussed in detail below. Despite entering the bloodstream, no detectable amount of dextran infiltrated the other organs. 1hr was determined to be the optimal infusion time as it minimizes off-target delivery while still covering the entire liver.

**Figure 3:**
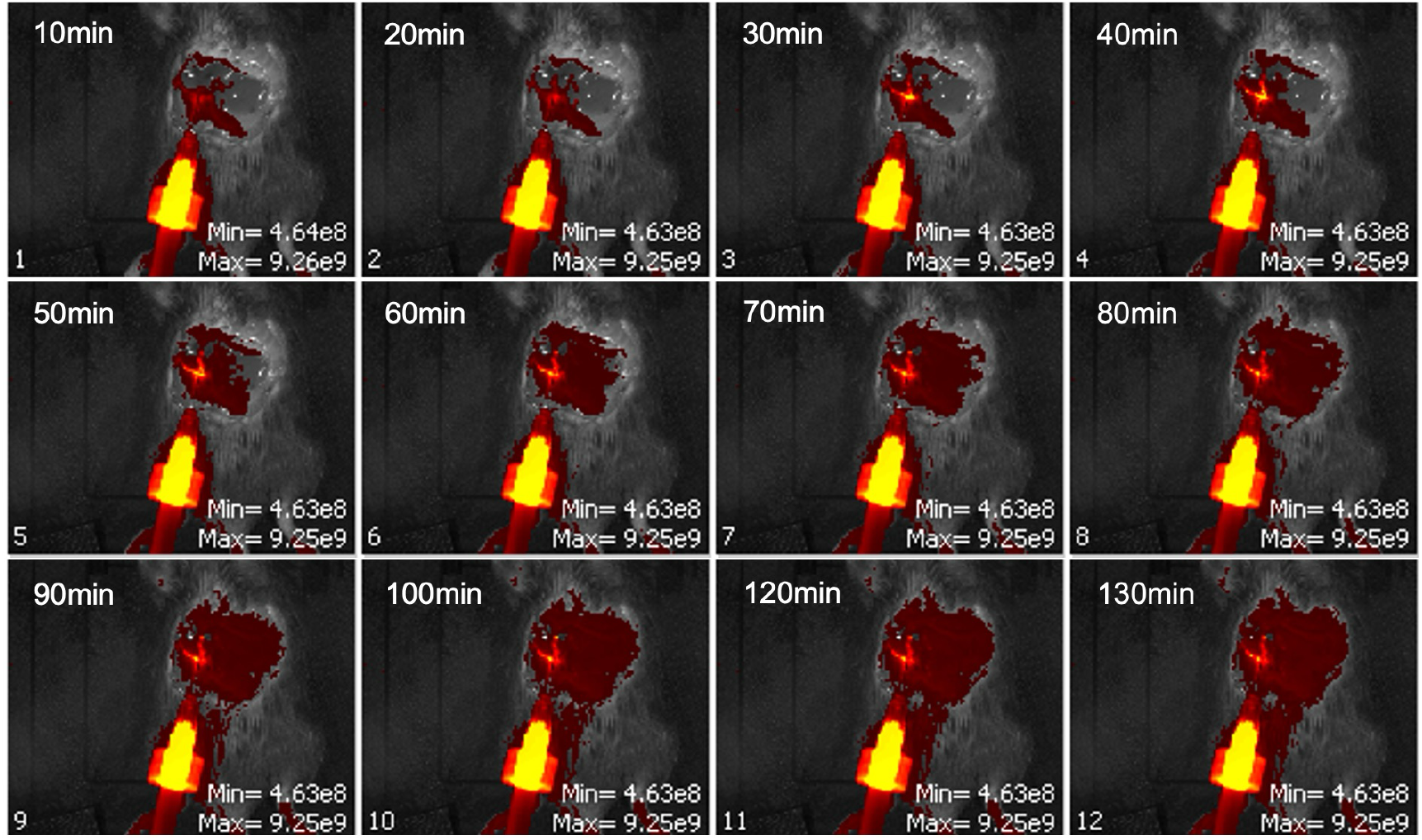
Live imaging of fluorescent dextran delivered via interstitial infusion of the liver. Fluorescent dextran is detected throughout the liver around 60mins of infusion. Beyond this time, the fluorescent signal in the liver is fairly consistent, while fluorescence of surrounding tissues gradually increases. Images acquired with IVIS Lumina Series III *in vivo* fluorescent imaging system, 0.5s exposure.

**Figure 4:**
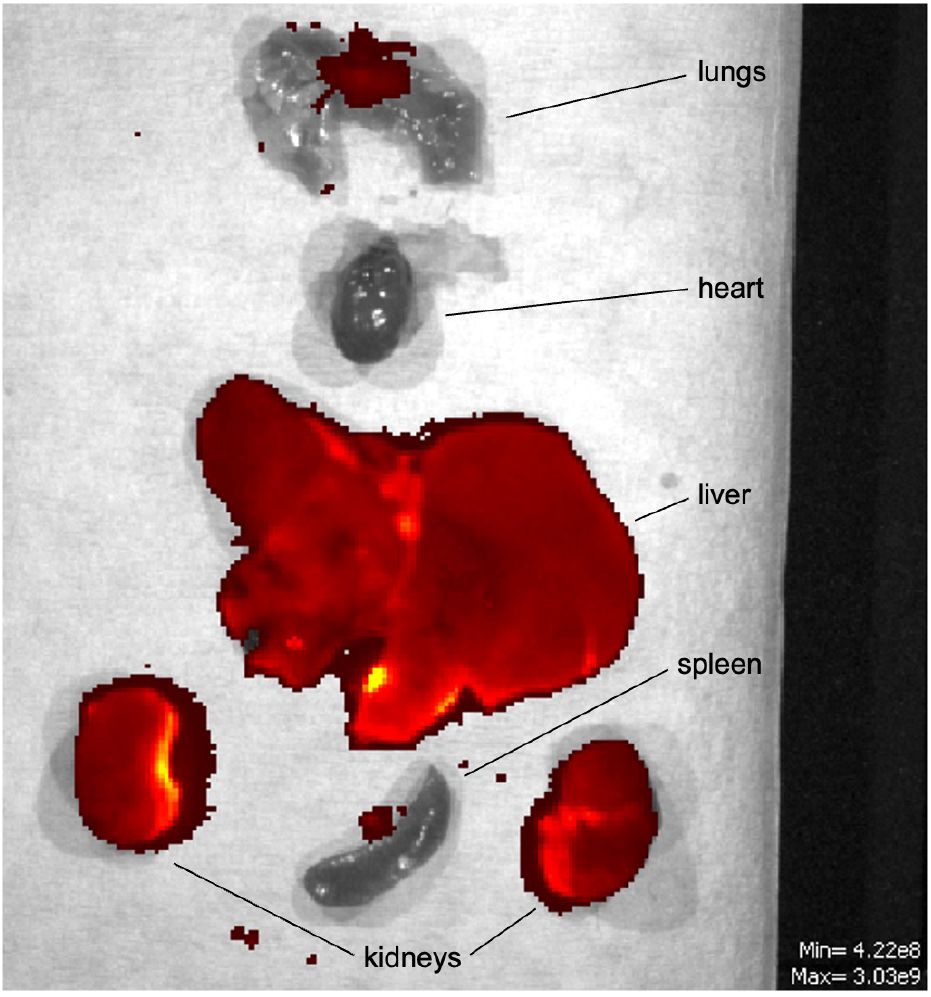
Whole organ imaging after interstitial infusion of fluorescent dextran in the liver for 60min. All lobes of the liver have a strong fluorescent signal, as well as both kidneys. The lungs, heart, and spleen show no significant signal. Image acquired with IVIS Lumina Series III *in vivo* fluorescent imaging system, 0.5s exposure.

Biodistribution after interstitial infusion in the liver was further characterized in mice up to 3 days after the infusion surgery. Fluorescence remained elevated and relatively stable in both the medial lobe of the liver, where the needle had been inserted, as well as the left lobe through 2 days after infusion (Fig. 5A,B). Similar results were seen in the renal cortex of the kidneys (Fig. 5C). By day 3, fluorescence of both organs was not significantly above the background level. Fluorescence of the lungs remained near background at all timepoints.

**Figure 5:**
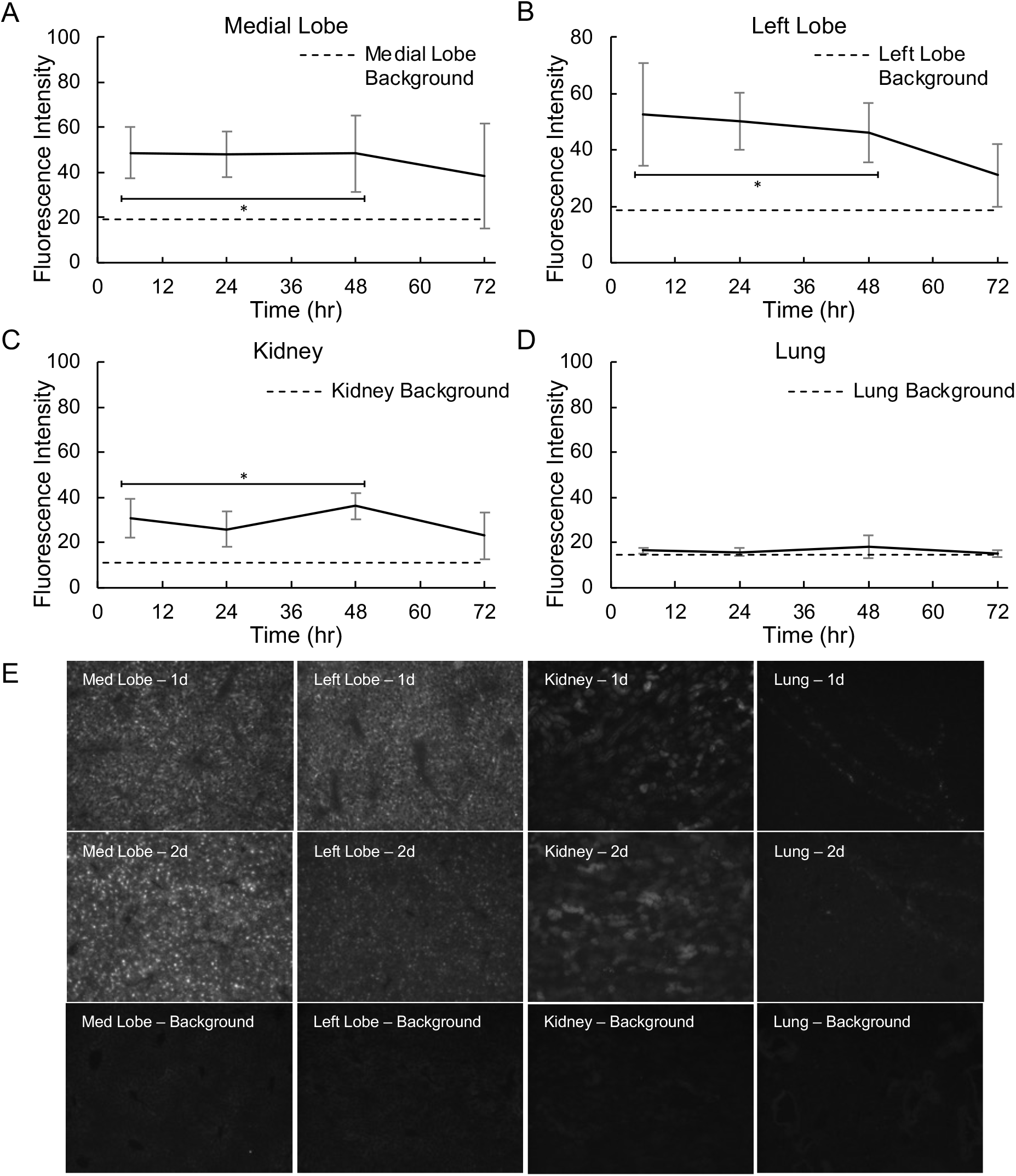
Quantified fluorescence after 60min interstitial infusion of fluorescent dextran in the liver. (A) Fluorescence in the medial lobe of the liver (the lobe where the needle was inserted) showed consistently elevated fluorescence through 2 days after infusion. By 3 days, fluorescence was not significantly different form background. (B) The left lobe of the liver, which was not directly infused, showed similar results. (C) Fluorescence was also detected in the renal cortex of the kidneys. This signal followed a similar trend to the liver, with stable elevation through 2 days. (D) Fluorescence of the lungs remained near background at all timepoints. (E) Representative fluorescence images taken at 10x magnification with Nikon Eclipse TE2000-S microscope. Plots depict mean ± SD; n=5 per timepoint. *p<0.05 compared to background controls (n=3). 750ms exposure time was used for liver and lungs. 75ms exposure time was used for kidneys due to higher fluorescence intensity.

### Purified CCH in vivo Toxicity & Efficacy

Preliminary *in vivo* tests were performed on mice maintained under anesthesia for up to 5hrs after infusion to determine an appropriate concentration of the purified CCH. Through these experiments, we found that concentrations greater than 1.25mg/mL were lethal. However, in our first experiments where anesthesia ended after the 1hr infusion period we found that mice that received concentrations of 0.25mg/mL or greater died shortly after withdrawal of anesthesia – despite these doses being viable in animals maintained under anesthesia for 5hrs after infusion until euthanasia. A concentration of 0.1mg/mL was chosen for the remaining experiments.

Based on the biodistribution data, potential risk targets for toxicity were the liver and kidneys. Serum markers of liver injury (AST, ALT) and kidney injury (creatinine) were measured at 1d and 5d after infusion (Fig. 6). AST and ALT were mildly elevated at 1d but fell close to or within normal range by 5d^23^. However, both markers showed similar elevations in mice infused with saline, suggesting this is related to liver injury from the procedure rather than the activity of the enzyme. There was no elevation in creatinine at any timepoint, indicating the enzyme does not cause acute kidney injury.

**Figure 6:**
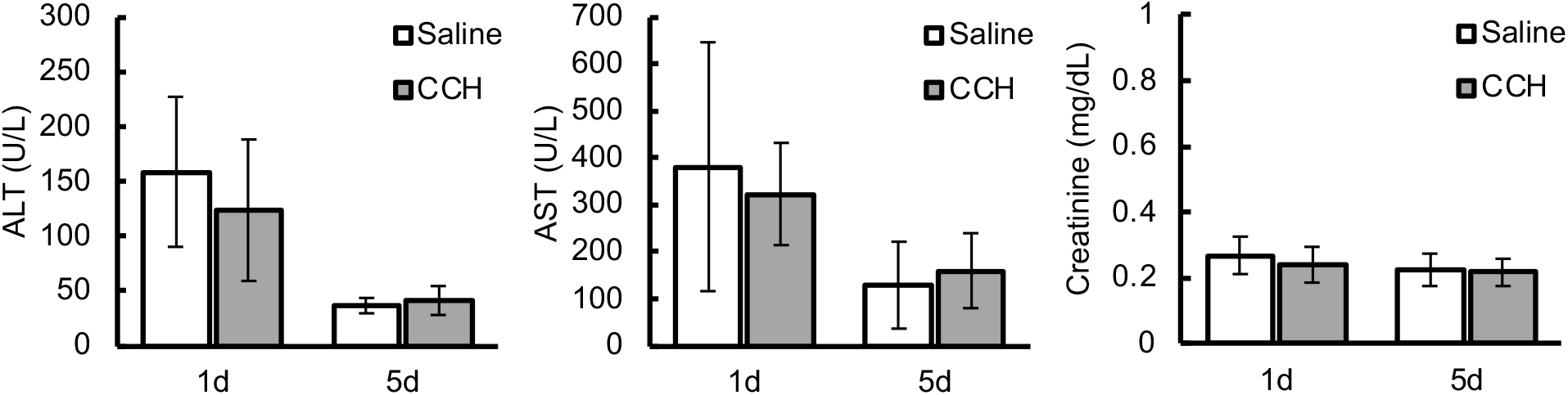
Serum injury markers at 1d and 5d after interstitial infusion of the cirrhotic liver with saline or purified CCH. Liver injury markers ALT and AST are elevated at 1d (reference range^23^: 20-82U/L ALT, 31-98U/L AST). ALT is normal by 5d, while AST remains slightly above the reference range. There is no significant difference between infusion with saline and purified CCH, suggesting these markers are not increased due to enzyme toxicity. Kidney injury marker creatinine is not elevated at any timepoint. Samples below the 0.2mg/dL level of detection for creatinine are not represented in that graph. Mean ± SD; 1d Saline n = 5, 1d CCH n = 5, 5d Saline n = 6, 5d CCH n = 6.

Treatment efficacy was assessed primarily by changes in fibrosis. Mice treated with purified CCH had a significant reduction in hepatic collagen at the 5d timepoint (Fig. 7A-C). Mean CPA was reduced by 38% in CCH-treated mice compared to saline controls.

**Figure 7:**
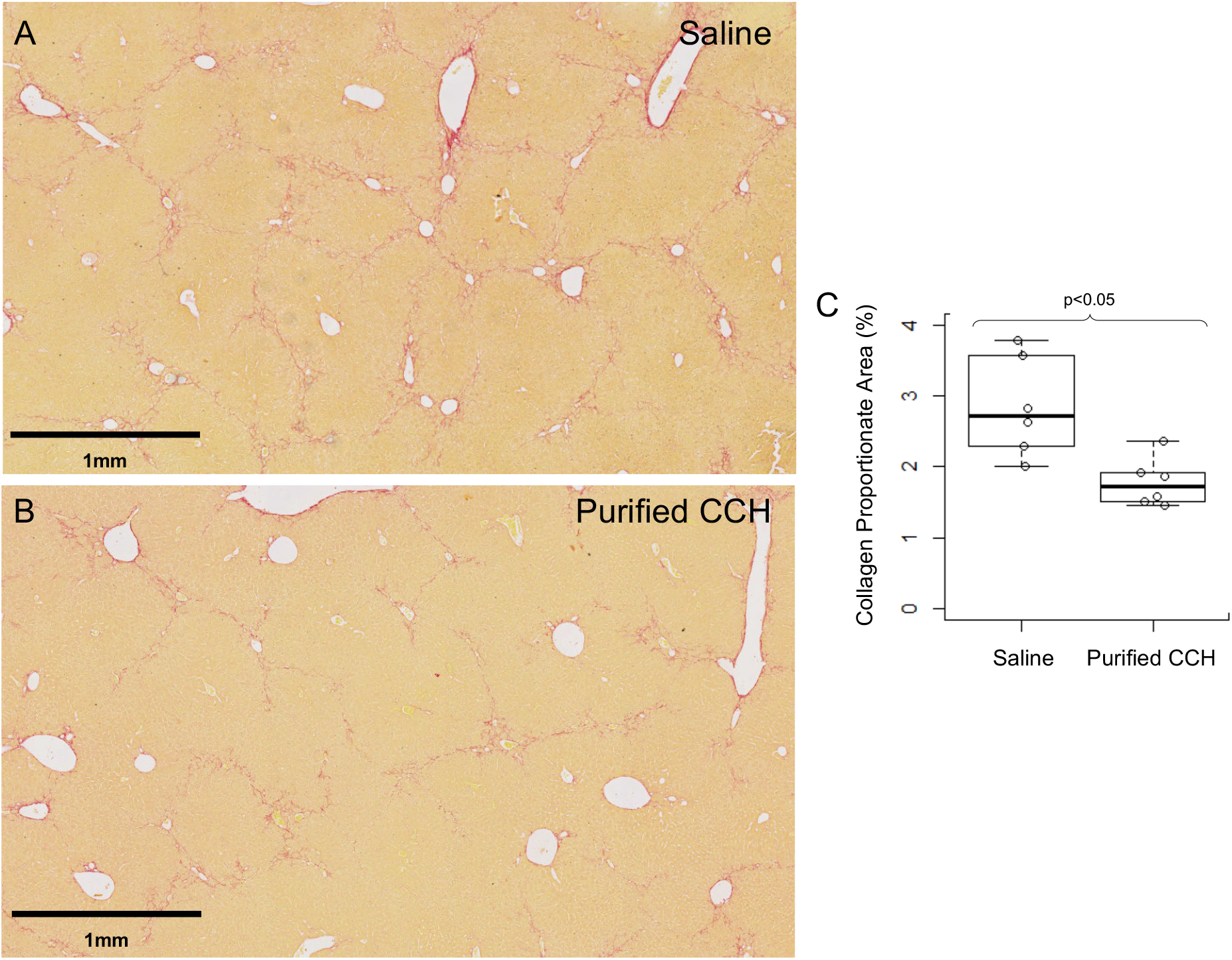
*In vivo* efficacy of interstitial infusion of purified CCH in the cirrhotic liver. (A,B) Representative picrosirius red staining 5d after infusion with saline or purified CCH, respectively. Fibrous bands in samples that received CCH are thinner and more disconnected. (C) Quantification of collagen in saline and CCH infused samples at 5d. Purified CCH caused a 38% decrease in collagen proportionate area.

## Discussion

Expression of matrix degrading enzymes is a common preclinical metric for prospective drugs for liver fibrosis, and there are also a number of papers describing gene therapies for various MMPs^24-27^. However, the delivery of an exogenous matrix-degrading enzyme has not been thoroughly investigated. Jin *et al*, 2005^28^ used a rabbit model of cirrhosis to perform biweekly injections of crude CCH through a portal vein catheter for 6-12 weeks. This study reported only 45% survival, which the authors attributed primarily to infection and surgical complications. However, it seems plausible that the presence of broad proteases in the infused enzyme mixture may have also contributed to the high mortality. Despite these complications, they were able to demonstrate significant histological improvements in the liver. El-Safy *et al*, 2020^29^ used a mouse model of moderate liver fibrosis to compare triweekly intravenous injection of crude CCH either free or coupled to chitosan nanoparticles that bind collagen. 100% of animals receiving free CCH died before the second dose, albeit with a sample size of 2, while 100% of animals receiving the optimal nanoparticle formulation completed the 3-week schedule. The authors reported moderate histological improvements in the CCH nanoparticle mice compared to controls. In both cases, a crude CCH mixture was used, and it was injected directly into the bloodstream. To our knowledge, ours is the first study to investigate direct interstitial infusion of collagenase in the liver.

Direct intrahepatic injection is a relatively common preclinical research tool used to seed cancer cells^30^ and stem cells^31^ in the liver. However, there is little prior work on the biodistribution of a macromolecule infused interstitially in the liver. Berraondo *et al*, 2006 tested intrahepatic injection for the delivery of an adeno-associated virus with a 50µL bolus injection^32^. They reported partial distribution within the injected lobe but not throughout the liver. Our results show that gradual infusion of 100µL reaches the entire liver, despite only directly infusing the medial lobe. This may simply be due to the difference in volume administered, but it is also likely that the gradual infusion improves interstitial distribution. Interstitial fluid drains through terminal lymphatics in response to small changes in interstitial pressure^33,34^. A bolus injection would cause a brief spike in local interstitial pressure around the injection site, allowing more of the injected material to drain away. An analogous effect has been reported in tumescent anesthesia, where gradual subcutaneous infusion of lidocaine dramatically reduces the plasma concentration^35^. The infusion protocol could further be refined in the future to be performed non-invasively under ultrasound guidance, rather than via laparotomy, to minimize impact on the recipient as well as enable investigation of multiple low-dose infusions over time. Repeated low-dose infusions may be clinically preferable so that the treatment can be titrated while monitoring liver function tests.

It is likely that there is also some distribution between lobes through the vasculature. The liver has a unique connection between intravascular and interstitial compartments due to the fenestrated sinusoidal endothelium. This connection deteriorates in advanced disease through capillarization of sinusoids, where fenestrations are lost and matrix accumulates in the space of Disse^36^. It is unclear how much of the connection remains in this model, and whether the more severe liver scarring seen in human disease may affect the biodistribution.

While in circulation, the structural collagen of the blood vessels would not typically be exposed to the CCH in the bloodstream due to the endothelial cell lining. In addition, early work with CCH demonstrated it is inhibited and cleared by alpha-2-macroglobulin, a large plasma protein that functions as a broad-spectrum protease inhibitor^37-39^. It is worth noting that alpha-2-macroglobulin is also produced by macrophages and hepatocytes, which may limit the activity of CCH within the liver^37,40,41^.

There are several notable limitations of our dextran experiments. First, there is a relatively small difference in size between the fluorescent dextran (70kDa) and purified CCH (116kDa). Both sizes are large enough that the primary distribution mechanism is convection^42^, so the short-term spread of these molecules is likely similar. The size differential is most relevant for the clearance through the kidneys. The fluorescent dextran is near the size threshold for glomerular filtration, though there is high variability based on shape and flexibility of the macromolecule^43,44^. The increased size of CCH likely reduces elimination through the kidneys compared to the fluorescent dextran. In addition, the dextran experiments do not account for specific removal processes for proteases and proteins in general. Overall, the dextran experiments primarily provide useful data for the short-term biodistribution of a macromolecule delivered via interstitial infusion in the liver, rather than an indication of their clearance and how long they persist in the tissue.

Interstitial delivery offers several advantages over intravenous routes previously investigated. Direct intrahepatic delivery ensures the highest concentration of the enzyme is at the desired site of action rather than in the plasma. Furthermore, as described above, the capillarization of sinusoids in advanced disease would impair infiltration of an enzyme injected in the bloodstream; with interstitial delivery, this would serve to maintain the enzyme concentration within the liver and limit systemic spread. Recent work has also suggested rethinking many interstitial structures previously believed to be densely-packed matrix – including the extrahepatic and intrahepatic biliary tree – as open fluid-filled spaces, which may have important implications for the spread of a macromolecule delivered via interstitial infusion^45^.

The results of our biodistribution and *in vivo* CCH experiments do not indicate any toxicity from the direct activity of the enzyme. The mild elevations in AST and ALT at 0.1mg/mL (matching elevations in saline controls), and the survival at concentrations as high as 1.25mg/mL when maintained under anesthesia suggest that the primary dose-limiting factor is not liver injury. It has been reported that certain anesthetic agents, including isoflurane, can be protective against sepsis and endotoxic shock. The mechanisms are not well understood, but isoflurane has been shown to enhance survival and delay systemic inflammation in murine models^46-50^. As we have shown a significant amount of the infused material enters the bloodstream, it is possible that contaminating endotoxin in the bacterial-origin CCH initiates a shock response which is inhibited until the animals are taken off anesthesia. If endotoxic shock is occurring, further purification could improve tolerance. As there appears to be no liver or kidney injury from the action of the enzyme, identifying and managing this reaction could enable higher doses and further improve treatment efficacy.

The functional deficiencies associated with cirrhosis can be attributed to several factors. Vascular changes leading to reduced perfusion are an important component^36,51-54^. However, there is also evidence to suggest scarring directly influences liver function. Hepatocytes in a stiff environment exhibit suppressed albumin production^55^, downregulation of cytochrome p450^56^ and HNF4a^57^ (two markers of hepatocyte differentiation and liver-specific function), as well as changes in a range of genes associated with normal epithelial function^58^. Hepatocyte senescence is also thought to contribute to liver dysfunction in cirrhosis^59^. The cause of this senescence is typically attributed to repetitive injury^60^; however, there is some evidence in age-associated senescence research demonstrating a link with tissue stiffness^61,62^, suggesting the local mechanical environment may also play a role in hepatocyte senescence. Therefore, improving the structural and mechanical aspects of cirrhosis may also be able to offer significant functional improvements.

It is well documented that mouse models of cirrhosis will naturally remodel and degrade the septa in the weeks after cessation of injury^63^. Importantly, this is not the case in humans. Cirrhosis in humans consists of much thicker and more stable fibrous septa which develop over many years. Research with the bile duct ligation model of fibrosis in rats showed that while spontaneous recovery was possible in this model, it required at least three times longer than it took to establish fibrosis^64^. This suggests that any prospective treatment for cirrhosis in humans will likely require highly aggressive collagen degradation to achieve any meaningful impact.

Longer term studies are needed to determine the functional consequences of the fibrosis reversal we have demonstrated. This is complicated by the spontaneous reversal discussed above. Rat models of cirrhosis develop thicker collagen septa from hepatotoxin exposure, and studies have shown long-term persistence of the septa after cessation of injury^17,65^. Confirmation of efficacy in the rat model will be an important next step.

We have described the biodistribution of a macromolecule delivered via interstitial infusion in the liver, and used these results to design an infusion protocol that covers the entire liver with minimal off-target delivery. We also used a mouse model of cirrhosis to show that purified CCH can rapidly reduce collagen content in the cirrhotic liver with only mild liver injury of the same level as saline infusion. Future work will investigate the long-term functional consequences of this treatment and investigate the mechanism of toxicity with a view to determining whether higher doses are feasible and beneficial. Development of a minimally invasive interstitial infusion procedure, potentially under ultrasound guidance, will also be investigated to determine the effect of multiple low-dose infusions over time.

