## Supplementary Figure for "Interstitial infusion of purified collagenase Clostridium histolyticum in the cirrhotic liver causes rapid reduction in fibrosis with minimal liver toxicity"

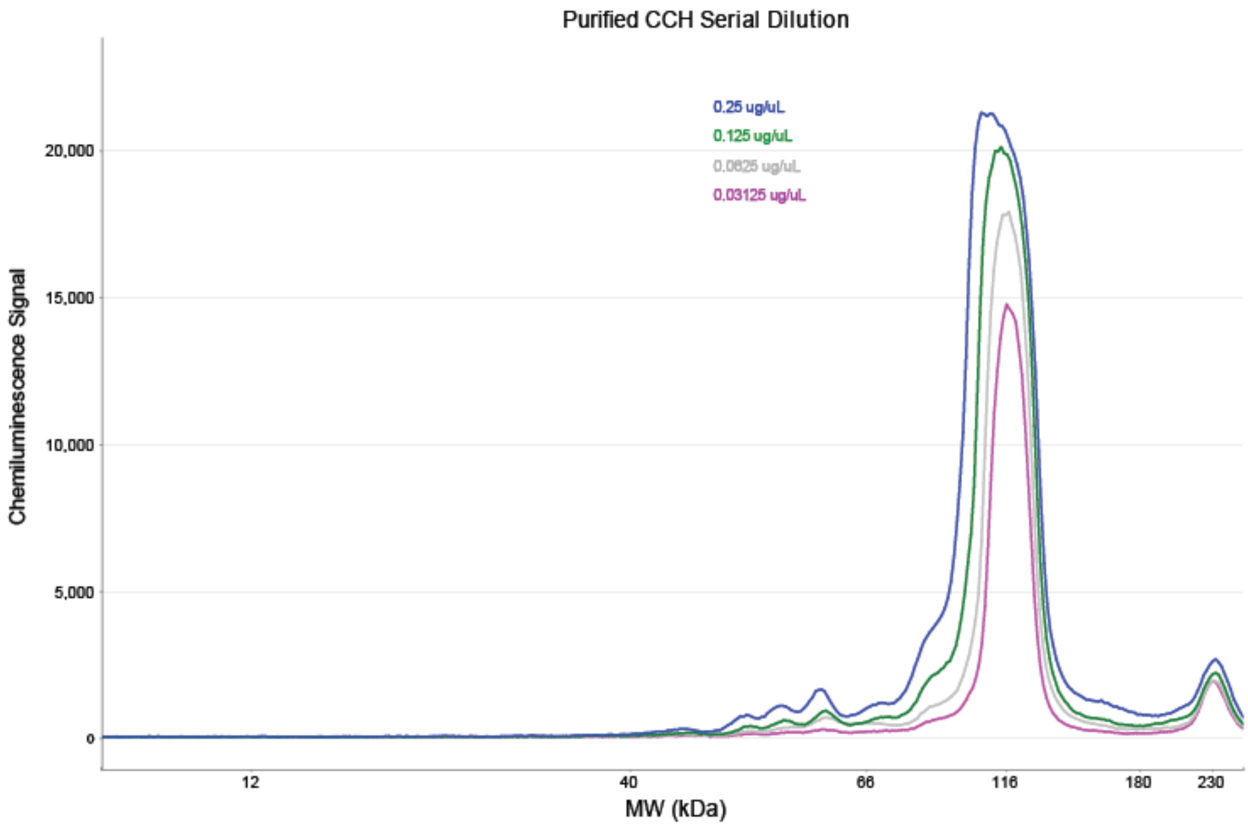


**Supplementary Figure 1: Molecular Weight Analysis for Serial Dilution of Purified CCH.** The small peak at 230kDa remains relatively consistent across the dilutions, suggesting it is likely an artifact rather than a component of the CCH.

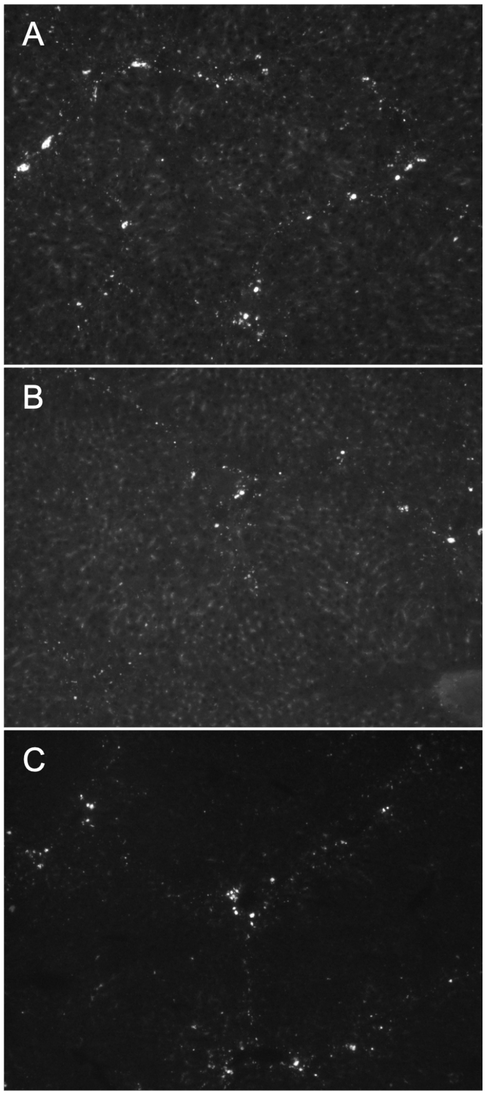


**Supplementary Figure 2: Interlobular distribution of fluorescent dextran tracer in the cirrhotic liver at 1d.** (A) Fluorescence in medial lobe (infused lobe). (B) Fluorescence in left lobe (non-infused lobe). (C) Background fluorescence in saline control. All images acquired at 10x magnification with Nikon Eclipse TE2000-S microscope.
